# TwinCell: Large Causal Cell Model for Reliable and Interpretable Therapeutic Target Prioritisation

**DOI:** 10.64898/2026.01.29.702072

**Authors:** Jean-Baptiste Morlot, Thomaz Dias, Sebastien Legare, Alessandro Romualdi, Elie Hatem, Yann Abraham

## Abstract

Drug discovery is impeded by the difficulty of translating targets from preclinical models to patients. In this work, we present TwinCell, a Large Causal Cell Model for target identification that, trained on *in vitro* cancer cell line perturbation data, generalises to patient-derived cell types while providing biologically meaningful interpretations of its predictions. Rather than predicting perturbation outcomes, TwinCell identifies the upstream regulators most likely to drive the transition between two cell states, such as diseased and healthy, by decomposing target probability over signalling paths through a multiomics interactome conditioned on single-cell foundation model embeddings. To validate this approach, we introduce TwinBench, a benchmarking framework that evaluates virtual cell models using recommendation system metrics while correcting for mode collapse through empirical *p*-value estimation. On both *in vitro* zero-shot scenarios and *in clinico* validation across five therapeutic areas, TwinCell outperforms not only state-of-the-art virtual cell models but also linear baselines and network-based methods, classically used to perform target identification. When applied to patient data, TwinCell recovers clinically approved drug targets and reconstructs known disease mechanisms, such as the type I interferon signalling cascade in Systemic Lupus Erythematosus, without any disease-specific supervision. These results demonstrate that constraining learned perturbation patterns to a biological interactome enables cross-tissue, cross-disease target identification with mechanistic interpretability, bridging the gap between high-throughput *in vitro* experiments and clinical insights.

## Introduction

Drug discovery remains a high-risk journey: only a small fraction of compounds ultimately reach patients, with estimates on the order of 10% progressing from early clinical trials to approval (1). This attrition is largely driven by the iterative nature of biological hypothesis generation involving time-consuming, costly experimental validations using *in vitro* and *in vivo* models. As a consequence, teams are routinely forced into high-stakes decisions under substantial uncertainty, committing to costly clinical validation before the confidence can compensate for the risks (2).

An analogy can be drawn from complex engineered systems. Historically, aerospace design relied heavily on iterative prototyping to identify failure modes and optimise architectures. However, this approach is limited by the impossibility of exhaustively testing the full parameter space (3). This paradigm shifted with the introduction of digital twins: high-fidelity digital counterparts of physical systems that support design exploration and validation prior to fabrication and deployment (3). Importantly, digital twins do not eliminate the need for physical validation; rather, they compress the expensive experimental iterations toward the end of the optimisation cycle while enabling broad *in silico* exploration early in development.

An essential value of a digital twin is its causal framework, providing interpretable and testable explanations of its behaviour. To extend this framework to natural systems, including biological ones, one must learn to identify the regulatory mechanisms governing cellular state transitions. Recent developments in world models have shown promising results in this direction. These models are composed of two components: a representation of the system’s accessible states and a model of its states’ evolution. Once trained, these systems can be used to guide an agent to navigate this latent space and identify the optimal perturbations that drive an initial state toward desired outcomes (4, 5).

Recent developments in machine learning have paved the way for digital twins of biological systems, termed virtual cells. Those models are trained to predict transitions between cellular states that are relevant to common biological questions, such as predicting the effect of a therapeutic intervention. Early approaches such as scGen (6) and CPA (7) use autoencoders to learn a latent space of the cellular state and predict perturbation effects across contexts. Graph-based methods like GEARS (8) leverage knowledge graphs to capture the biological relationships between genes, improving generalisation to unseen perturbations by explicitly modelling gene-gene dependencies. Meanwhile, the growing availability of large single-cell atlases and the success of foundational models in other domains results in the emergence of new large-scale pretrained embeddings such as scGPT (9) and Geneformer (10), which learn biologically meaningful gene representations from millions of cells. Finally, large-scale perturbation datasets including scPerturb (11) and Tahoe-100M (12) now provide the data required to train models that can predict transcriptomic changes across diverse cellular contexts.

The Arc Institute’s recent STATE model (13) exemplifies this emerging class of models: trained on multiple large scale perturbation databases, including Tahoe-100M, it predicts transcriptomic changes due to perturbations across diverse contexts by learning to navigate into a pretrained latent space. Starting from an unperturbed cellular state and a perturbation, the STATE model predicts the perturbed cellular state, enabling downstream exploratory tasks. However, generalising perturbation effects to entirely new cell types or novel perturbations remains a challenge, as recent work demonstrated that linear models can outperform deep learning approaches when extrapolating to unseen contexts (14).

This finding underscores both the need for challenging, biologically meaningful benchmarks and the importance of developing interpretable models that capture causal relationships rather than optimising global transcriptional similarity. Conventional benchmarks relying on metrics such as Pearson correlation or mean squared error can be misleading in high-dimensional transcriptomic spaces: they may reward average-case predictions, fail to detect mode collapse, and overlook intervention-specific signatures (13, 15–17).

In this work, we address a complementary formulation of the perturbation modelling problem by extending the approach introduced in PDGrapher (18). Rather than predicting perturbation outcomes from an initial state, we frame the task as a **perturbation recommendation system**: given an initial state and a desired target state, we want to predict the perturbations that most likely drive the transition between them. This formulation aligns directly with the practical needs of drug discovery, where the goal is to identify interventions that reverse disease phenotypes.

We introduce TwinCell, a Large Causal Cell Model (LCCM) that computes the probability of target candidates given observed Differentially Expressed Genes (DEGs) and cell-state context, decomposing this probability over signalling paths through a multiomics interactome conditioned on foundation model embeddings. Trained on a single *in vitro* perturbation database (Tahoe-100M (12)), we evaluate TwinCell’s generalisation capability both *in vitro* (zero-shot cell lines and perturbations) and *in clinico* across five therapeutic areas.

We show that TwinCell outperforms not only state-of-the-art virtual cell models but also linear baselines and network-based methods classically used to perform target identification. Applied to Systemic Lupus Erythematosus, the model recovers clinically approved drug targets, recapitulates the canonical type I interferon signalling cascade, and identifies *de novo* target candidates supported by interpretable causal paths. These results demonstrate that constraining learned perturbation patterns to a biological interactome enables cross-tissue, cross-disease target identification with mechanistic interpretability.

## Results

### TwinCell: Large Causal Cell Model for interpretable target identification

TwinCell is a scalable probabilistic causal framework for target identification (Figure 1a). Given two cell states, for example a diseased and a healthy condition, TwinCell identifies the most likely upstream regulator *t*^∗^ driving the observed transcriptional differences between them:

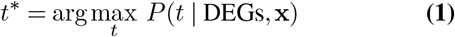

where DEGs are the differentially expressed genes between the two conditions and **x** = (**x**_1_, **x**_2_) are cell-state embeddings obtained from the Geneformer foundation model (10). Assuming that signalling paths to different DEGs propagate independently through the interactome, TwinCell decomposes this probability as:

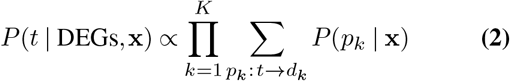

where each *P* (*p*_*k*_ | **x**) is the probability of a signalling path from candidate target *t* to DEG *d*_*k*_ (see Material & Methods). The model learns cause–effect relationships by training end-to-end on *in vitro* interventional perturbation data, using a multiomics interactome as an inductive bias to constrain signal propagation to peer-reviewed molecular interactions, and cell-state embeddings from the Geneformer foundation model (10) to learn the mapping from cellular context to edge weights. Once trained, TwinCell can prioritise candidate targets and construct causal graphs for any pair of cell states, providing mechanistic hypotheses for experimental follow-up.

**Fig. 1.**
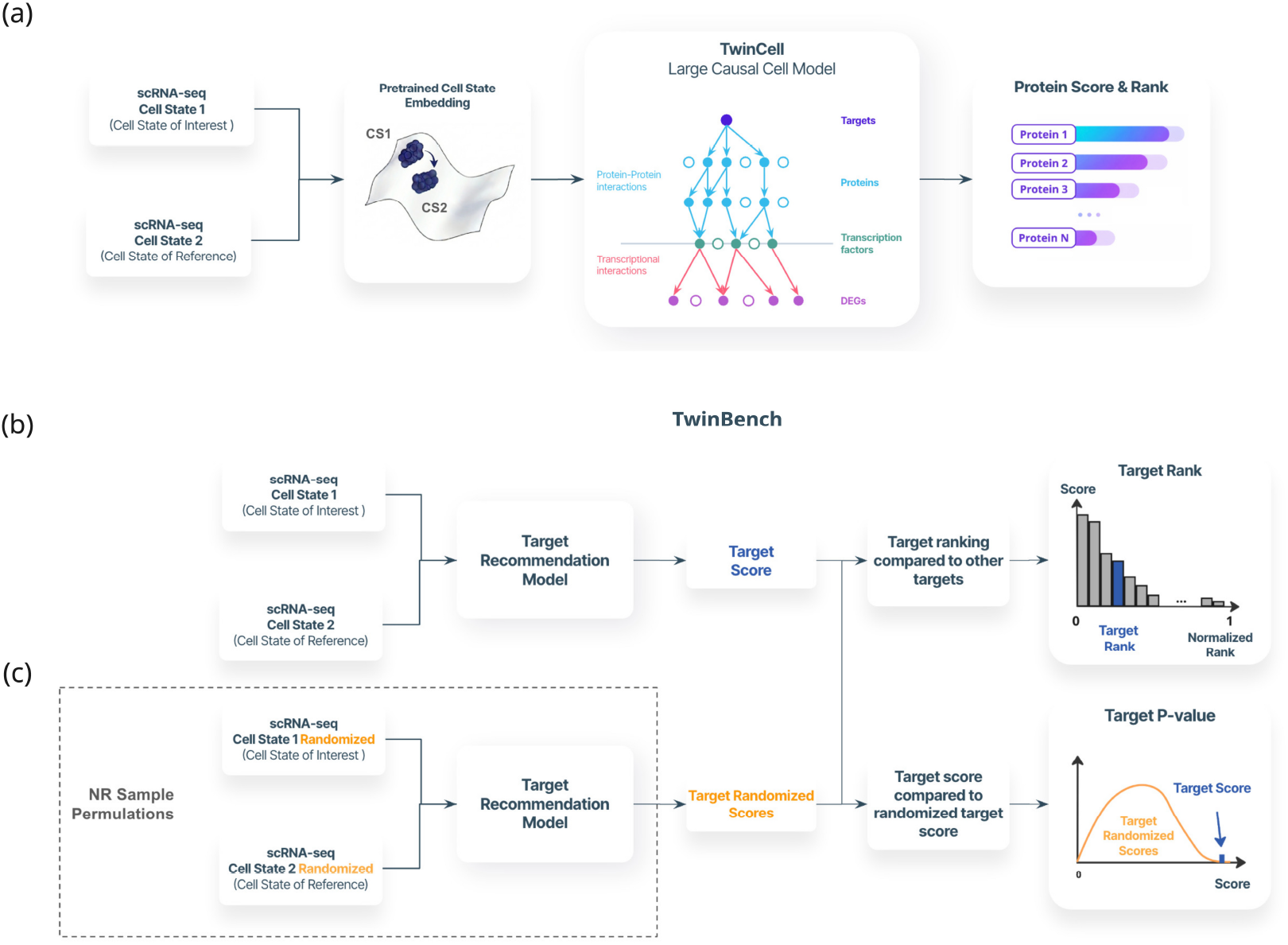
TwinCell Large Scale Causal Cell Model architecture and TwinBench evaluation framework for virtual cell models. (a) TwinCell model architecture. We select two cell states, the state of reference as starting point and the state of interest as objective, using single cell RNA-seq data. Each sample is embedded using a pretrained foundation model, here Geneformer (10), and used to generate a cell state specific causal graph, that estimates the probability of each protein as candidate target given the observed differentially expressed genes (DEGs). The model uses a multiomics interactome as inductive prior combining transcriptional regulatory and protein–protein interactions: *P* (Target | DEGs, x_1_, x_2_). (b) TwinBench evaluation pipeline for virtual cell models using recommendation system metrics. First the model predicts a score for each target given a pair of cell states. Considering the score of the true target, we can compute its rank by comparing it to the scores of the other targets, the higher the score, the lower the rank. (c) Then we recompute the score by permuting the input data, and calculate the fraction of permutations in which the permuted score exceeds the actual score. This empirical *p*-value quantifies model uncertainty and corrects for models that predict the same targets regardless of input signal, aka popularity bias in recommendation systems or mode collapse in generative models.

### TwinBench: Robust benchmarking framework for virtual cell models

Current evaluations of virtual cell models rely on similarity metrics (e.g. Pearson correlation, MSE) between predicted and observed transcriptomic profiles. These measures can mask a common failure mode known as mode collapse in generative models or popularity bias in recommendation systems: the model reproduces the training distribution’s most frequent outcomes while ignoring the specific input signal (13, 14). Such predictions appear accurate because high-dimensional expression profiles share substantial baseline correlation, but carry no perturbation-specific information. By recasting evaluation as a recommendation problem, where the model must rank the correct target above thousands of alternatives, the benchmark connects directly to target identification workflows and exposes input-insensitive predictions that similarity metrics miss.

We introduce TwinBench, a benchmarking framework that evaluates the performance of virtual cell models in a recommendation system setting, while correcting for popularity bias or mode collapse (Figure 1(b-c)). In conventional recommendation settings, recall and precision are computed at a fixed rank cutoff *K*, selecting the top-*K* items and evaluating how many true positives are retrieved (Figure 1b). However, out-of-distribution evaluation requires additional care: even when presented with random noise, recommendation models do not predict uniformly at random but instead favour items frequently observed during training—a phenomenon known as popularity bias. To address this, we replace the fixed *K* threshold with an empirical *p*-value computed independently for each gene. Specifically, we assess each gene’s score against a null distribution obtained by permuting input data, summing how often the permuted score exceeds the observed score (Figure 1c). The empirical *p*-value for target *t*_*j*_ is defined as:

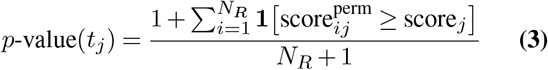

where *N*_*R*_ is the number of random permutations. This formulation establishes a gene-specific significance threshold: targets that consistently achieve high scores regardless of input signal (due to interactome topology or training frequency) will not reach significance, whereas targets whose scores depend on the input differential expression will.

To estimate the model performance, we summarise the results across multiple sample pairs and targets into normalised rank and empirical *p*-values distributions. From these distributions, we can derive two metrics to summarise the model performance parametrised by p-value threshold *pv*: the Inverse Mean Normalised Rank (IMNR), defined as the inverse of the mean normalised rank of true targets with *p*-value *< pv* and the recall, defined as the fraction of targets with *p*-value *< pv*. We combine these scores into the F1-score across different *p*-value thresholds and derive the area under the F1-score curve (AUC F1-score). Given a *p*-value threshold *pv*:

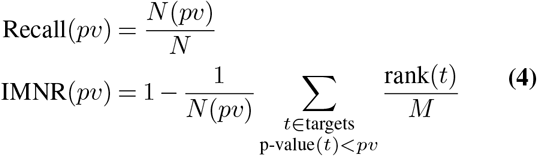

with

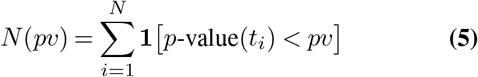

the number of true targets with *p*-value below *pv, N* the total number of true targets, and *M* the number of genes expressed in at least one of the two cell states. Normalising the rank by *M* rather than by the total number of candidate genes ensures that unexpressed genes do not inflate the score. In this work, we sample 30 linearly-spaced *p*-value thresholds in [0.01, 1.0] Figure 1c. Next we investigate TwinCell generalisation capabilities *in vitro* and *in clinico*.

### TwinCell outperforms state-of-the-art models on *in vitro* out-of-distribution generalisation scenarios

We benchmarked TwinCell against five baselines (Figure 2b): two linear Ridge regression models adapted from EASE (19), a simple yet competitive recommendation-system baseline, trained on either highly variable genes (HVGs) or Gene-former embeddings; a random baseline using shuffled HVG features (Random HVG); Network Medicine (20), a leading drug-repurposing method based on network proximity; and the STATE model (13), a state-of-the-art virtual cell model. The Random HVG baseline feeds randomly shuffled gene expression vectors to the HVG Ridge regression model, destroying all biological signals while preserving the marginal distribution. While TwinCell and the linear models explicitly predict perturbation targets, Network Medicine and STATE rank drugs. Because TwinBench evaluates recommendation quality and penalises popularity bias, it is agnostic to whether the ranking applies to targets or drugs, provided the ground truth is defined accordingly (see Material & Methods).

**Fig. 2.**
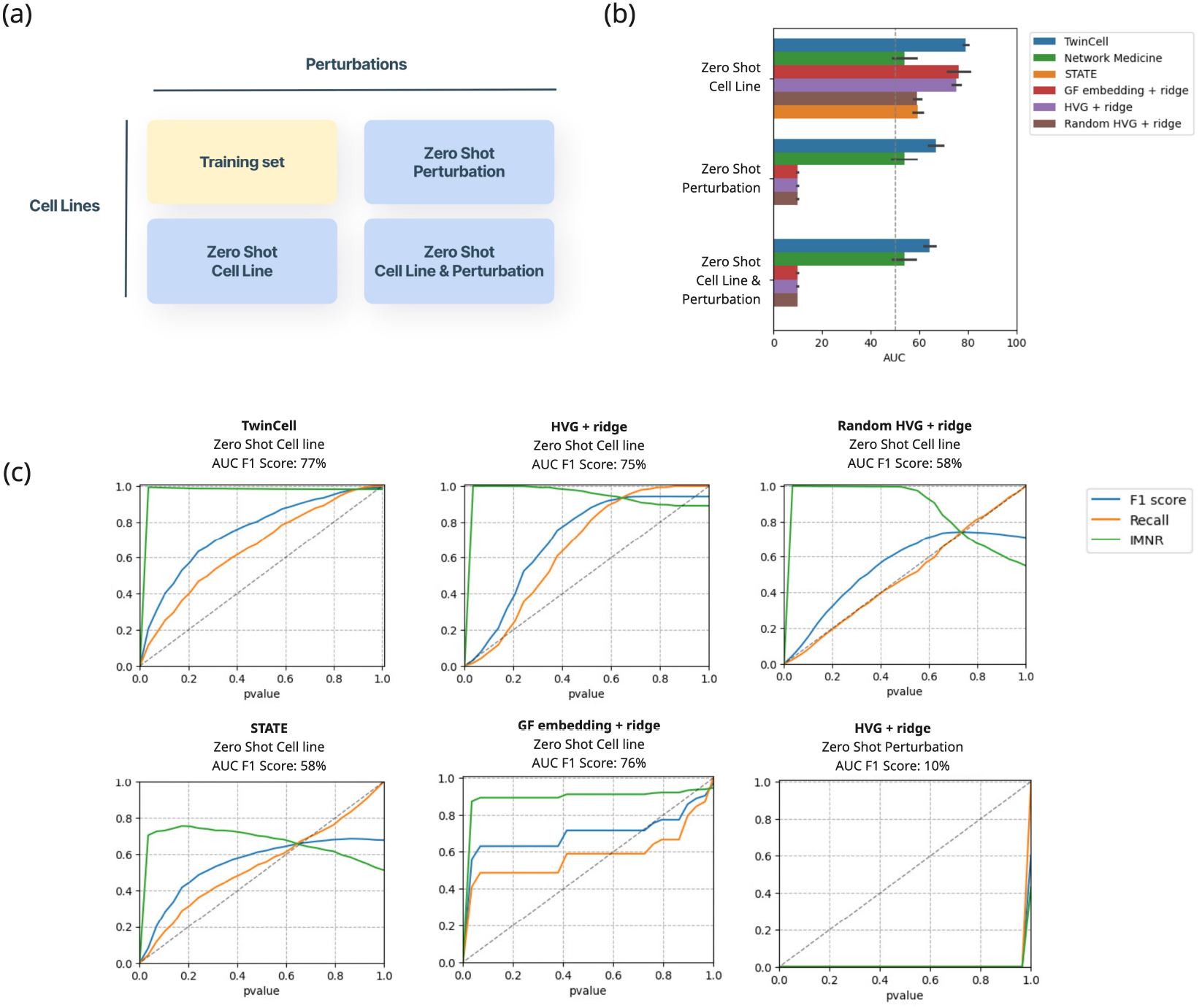
*In vitro* generalisation scenarios and model comparison. (a) Zero-shot generalisation scenarios corresponding to held-out cell lines, held-out perturbations, or both. (b) Model performance across generalisation scenarios measured by TwinBench AUC F1 score using drug targets as ground truth. The STATE model (13) is shown only for the zero-shot cell line scenario, as it requires the perturbation identity as input and therefore cannot be evaluated on unseen perturbations and Network Medicine is the only model whose performance is agnostic of the training set. (c) Decomposition of the AUC F1 score across *p*-value thresholds for the zero-shot cell line scenario, showing IMNR, Recall and F1 curves per model and one example of zero-shot perturbation scenario for linear baselines.

We evaluated all models across three zero-shot generalisation scenarios (Figure 2a): Zero-shot cell lines, zero-shot perturbations, and zero-shot cell lines and perturbations simultaneously (list available in Tables S1 and S2). Across all scenarios, TwinCell consistently outperforms the other methods (Figure 2b). In the zero-shot cell line scenario, where a direct comparison with STATE is possible, TwinCell achieves the highest AUC F1 score, while STATE performs on par with the Random HVG baseline. Network Medicine, whose performance is independent of training data but remains sensitive to DEG quality, performs near the random baseline. In the zero-shot perturbation and combined scenarios, TwinCell maintains strong performance. In contrast, the linear baselines fail to generalise to unseen perturbations, translating into a *p*-values close to 1 (Figure S1a), and an AUC F1 down close to ∼10%.

To illustrate how TwinBench discriminates between models, we decompose the AUC F1 score into its IMNR, Recall and F1 components across *p*-value thresholds for the zero-shot cell line (Figure 2c). TwinCell maintains high IMNR and Recall across thresholds, indicating that its top-ranked predictions are both accurate and input-specific. The Random HVG baseline serves as a useful control: it exhibits the characteristic signature of popularity bias, with high IMNR but Recall close to the diagonal, confirming that predictions are insensitive to the input signal. STATE shows a similar pattern, with Recall near the diagonal despite high IMNR. Next we investigated TwinCell’s performance in identifying disease-associated pathways.

### TwinCell recovers clinically validated drug targets with interpretable causal evidence in Systemic Lupus Erythematosus

We tested TwinCell on a first *in clinico* example using a dataset from Perez et al. (21). Blood samples were collected from healthy donors and patients with Systemic Lupus Erythematosus (SLE) (Figure 3a).

**Fig. 3.**
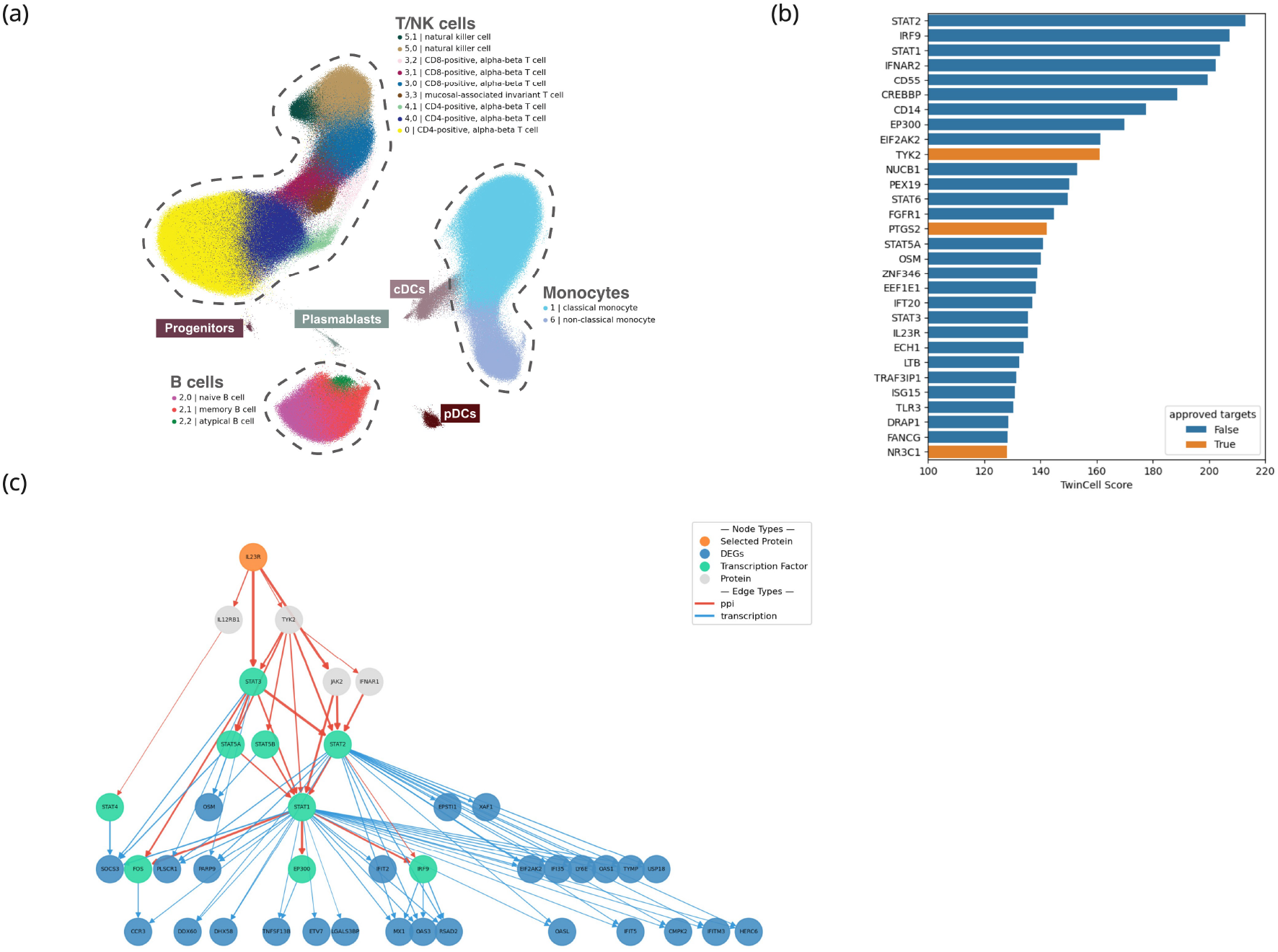
Causal analysis of Systemic Lupus Erythematosus in activated CD4^+^ T cells. (a) UMAP visualisation of the SLE atlas with cell-type annotation. (b) Top 30 proteins ranked by TwinCell score in activated CD4^+^ T cells (cluster 4, 1 in (a)). The score represents the log-unnormalised posterior log *P* (*t* | DEGs, x) up to an additive constant shared across all targets. Approved targets from OpenTargets (phase III/IV clinical trials) are highlighted in orange. (c) Causal graph representing the inferred top 5 most probable signalling paths from IL23R receptor in activated CD4^+^ T cells to DEGs. For visual clarity, only DEGs *d*_*k*_ with a probability exceeding 20-fold enrichment over the uniform baseline are shown, i.e. *P* (*d*_*k*_ | *t*, x) *>* 20*/M*, with *M* is the number of genes expressed in at least one of the two cell states. Orange nodes correspond to the selected protein (IL23R), green nodes represent Transcription Factors, blue nodes correspond to DEGs, and grey nodes to other intermediate proteins. Red edges correspond to protein–protein interactions and blue edges to protein–gene (transcriptional) interactions. Edge thickness reflects the cell-state-specific weights learned by the model, with thicker edges indicating more probable routes.

After re-annotating the cell types, we focused our analysis on CD4^+^ T cells, a disease-relevant population in which inflammatory signalling programmes are directly actionable by clinically established immunomodulatory targets (22). We further focused on activated CD4^+^ T cells (cluster 4, 1 in Figure 3a) as it represents a distinct interferon-imprinted activation state within the CD4 compartment, characterized by elevated ISG and cytoskeletal remodeling gene expression, making it a biologically relevant, interferon-driven CD4^+^ T cell state in SLE. We computed Differentially Expressed Genes (DEGs) comparing activated CD4^+^ T cells from SLE patients to those from healthy donors and used these DEGs as input to the model. Each protein in the interactome is ranked by the TwinCell score, and the top 30 proteins are shown in Table S4 and Figure 3b. To assess whether TwinCell recovers clinically validated biology in this setting, we obtained the list of approved drugs for SLE from OpenTargets, focusing on drugs in phase III and IV clinical trials. In total, from all approved targets expressed in CD4^+^ T cells (27 targets), 45% (12 targets) are ranked within the top 5% of TwinCell’s predictions (Table S5). Notably, the ranking highlights multiple clinically established axes of SLE immunomodulation, including the interferon receptor pathway (IFNAR1) and downstream JAK/STAT signalling components (TYK2, JAK1/2/3), alongside additional validated targets such as PTGS2 and NR3C1 (Figure 3b). Beyond approved clinical targets, we decided to focus on IL23R (rank #21), the receptor for the IL-23 cytokine (Figure 3c). The IL-23 axis is a central pathway in autoimmunity, and therapeutic blockade has demonstrated clinical utility across multiple inflammatory diseases, including psoriasis, Crohn’s disease and ulcerative colitis (23–25). TwinCell also provides mechanistic interpretability by decomposing target probabilities over signalling paths in the interactome. As an exemple, TwinCell reconstructs causal paths linking IL23R through JAK/STAT intermediaries to TNFRSF13B (BAFF), a cytokine known to be involved in the disease (26) (Table S6, Figure 3c).

The resulting causal subgraph provides an interpretable rationale for the IL23R prediction, demonstrating how TwinCell jointly supports target prioritisation and explainability through causal reasoning constrained by biological network structure. These results were obtained without any SLE-specific training, illustrating TwinCell’s ability to transfer mechanistic knowledge learned from *in vitro* perturbation experiments to recover clinically validated targets in a disease- and cell-type-specific manner. Next we tested TwinCell performance across multiple diseases.

### TwinCell outperforms target identification and drug repurposing methods in the retrieval of clinically validated targets in multiple independent diseases

Next, we compared TwinCell to other target identification methods on patient-derived data. We applied the TwinBench framework to five diseases spanning distinct tissues and pathologies, all unrelated to the cancer cell lines used during training: Ulcerative Colitis and Crohn’s Disease (intestine) (27), Systemic Lupus Erythematosus (SLE, blood), Parkinson Disease (brain), and Psoriasis (skin). For each disease, we selected a cell type with established pathogenic relevance: enterocytes in Ulcerative Colitis and Crohn’s Disease, which form the intestinal epithelial barrier whose dysfunction is central to inflammatory bowel disease (27); activated CD4^+^ T cells, a subset associated with disease activity (22); oligodendrocytes in Parkinson Disease, recently implicated in neurodegeneration through transcriptomic studies of the substantia nigra (28); and Th17 cells in Psoriasis, the key effector population driving IL-17-mediated skin inflammation (29). Disease-specific approved targets were extracted from OpenTargets (phase III/IV clinical trials). To assess the models’ ability to generalise beyond targets seen during training, we compared performance across three scenarios: all targets (Figure 4a), in-train targets only (Figure 4b), and out-of-train targets only (Figure 4c). In each graph the vertical dotted line corresponds to random performance. The full list of targets per disease and scenario is available in Table S3. Network Medicine serves as a reference throughout, since its performance is independent of the training set.

**Fig. 4.**
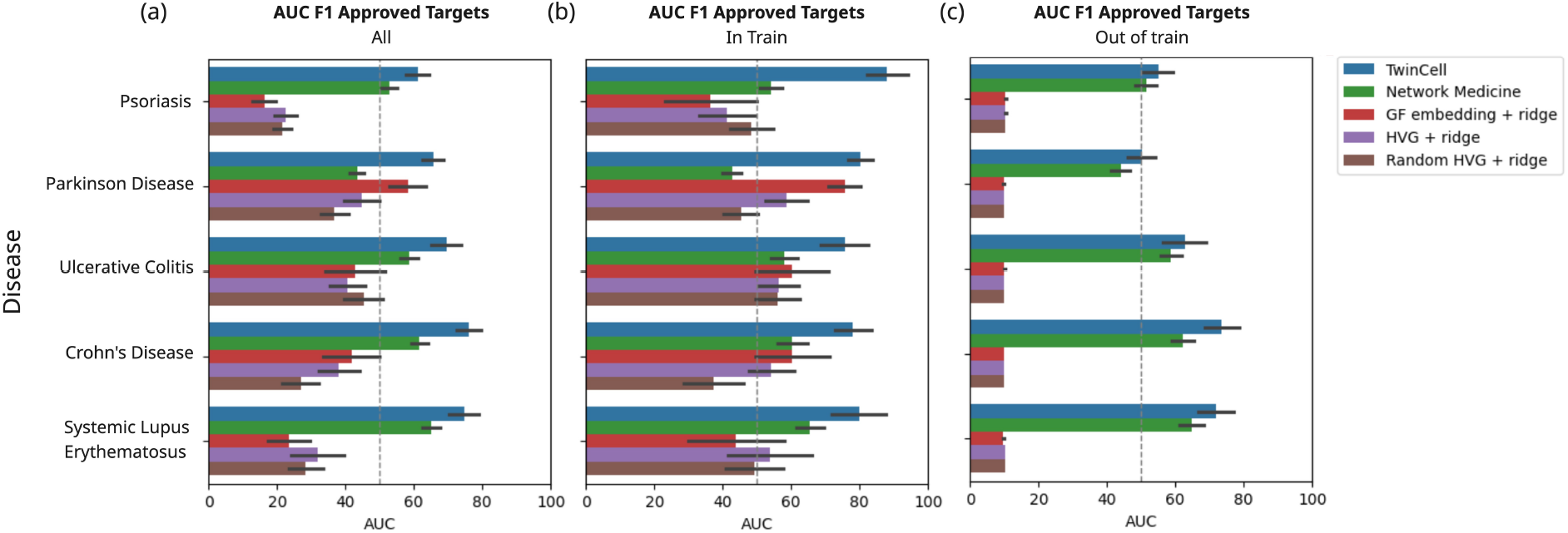
*In clinico* scenario results across five therapeutic areas. (a) TwinBench AUC F1 scores for four models across five diseases, using approved targets from OpenTargets as ground truth: TwinCell (blue), Network Medicine (green), and two Ridge regression baselines using Geneformer embeddings (red) and Highly Variable Genes (purple) as input. Dark segments correspond to 95% confidence intervals. For each disease, results are shown for a disease-relevant cell type: Th17 cells in Psoriasis, oligodendrocytes in Parkinson Disease, enterocytes in Ulcerative Colitis and Crohn’s Disease, and CD4^+^ T cells in SLE. (b) Same as (a) restricted to targets present in the training set. (c) Same as (a) restricted to targets absent from the training set.

Across all three scenarios, TwinCell outperforms Network Medicine and linear baselines (Figure 4a-c). In the all-targets scenario (Figure 4a), TwinCell consistently achieves the highest AUC F1 scores across all five diseases. The advantage over Network Medicine is most pronounced in the in-train scenario (Figure 4b), where TwinCell can leverage the mechanistic patterns learned from perturbation data to complement the interactome topology. In the out-of-train scenario (Figure 4c), both TwinCell and Network Medicine outperform linear methods, confirming that using an interactome as a structural prior is critical for generalisation to novel targets. Accordingly, linear baselines largely fail when targets fall outside their training distribution, yielding high *p*-values and low AUC F1 scores (Figure S1a).

Taken together, these results demonstrate TwinCell’s ability to generalise from *in vitro* cell line perturbation data to *in vivo* disease-relevant cell types, recovering clinically validated drug targets across five independent therapeutic areas.

## Discussion

We present TwinCell, a scalable probabilistic framework for interpretable target identification that learns cell-state-specific causal graphs over a large-scale multiomics interactome. Rather than predicting perturbation outcomes, TwinCell focuses on the inverse problem: given two cell states, identify the upstream regulators driving the transition between them. This framing aligns more directly with the practical needs of drug discovery, where the goal is to identify interventions that reverse disease states. By integrating single-cell foundation model embeddings with an interactome, TwinCell decomposes target probability over signal transduction paths, constraining predictions to mechanistically plausible routes. Trained on Tahoe-100M *in vitro* cancer cell line perturbation data, the model generalises to patient-derived cell types across five therapeutic areas, recovering clinically approved drug targets supported by causal graphs that link targets to DEGs and highlight the biological rationale behind each prediction.

A core value of virtual cell models lies in their ability to generalise to unseen contexts, both to reduce the experimental burden and to generate actionable insights before committing to clinical trials. To rigorously assess this capability, we designed a benchmarking framework that evaluates models across two complementary scenarios: *in vitro* zero-shot generalisation, where held-out cell lines, perturbations, or both are withheld from training; and *in clinico* retrospective analysis, where models are tasked with recovering clinically approved drug targets across five therapeutic areas and disease-relevant cell types. We benchmarked TwinCell against five baselines across four methodological families (Figure 2b): two linear Ridge regression models adapted from EASE (19), a simple yet competitive recommendation-system baseline, trained on either Highly Variable Genes (HVGs), a common gene selection method or Geneformer embeddings (10), a well established foundation model; (ii) one network-based method, Network Medicine (20), a leading drug-repurposing approach based on network proximity; (iii) a state-of-the-art virtual cell model, STATE (13) and (iv) a random model using shuffled HVG features (Random HVG) serving as reference. Across all *in vitro* zero-shot scenarios, TwinCell out-performed every baseline, with its strongest advantage on held-out perturbations, that directly tests generalisation to novel targets. In the *in clinico* scenario, TwinCell consistently recovered approved drug targets across all five therapeutic areas. While performance was expectedly higher for targets present in the training set, TwinCell also recovered approved targets absent from training, demonstrating its suitability for *de novo* target identification and drug repurposing. The comparison with Network Medicine and Linear model with Foundation Model embeddings highlights the importance of learned perturbation patterns over interactome constraints. TwinCell’s performance suggests that neither the interactome alone nor perturbation data is sufficient, it is their combination that enables generalisation to novel targets in unseen cell types. This finding suggests that the path to reliable virtual cell models lies not in ever-larger datasets alone (14), but in effectively integrating learned representations with high-quality biological knowledge. Future evaluations should also increase the size of the cell lines and perturbation landscape to hold out entire tissues or pathway classes rather than individual cell lines or compounds, building on the strong cross-tissue evidence already provided by the *in clinico* results.

Focusing on SLE, we show that the causal graph captures the relations between DEGs and IFN-*α* in activated CD4^+^ T cells, recapitulating known disease biology. This suggests that causal patterns learned from *in vitro* cancer cell line perturbations encode transferable regulatory principles that generalise across tissues and disease contexts. Beyond recovering known mechanisms, TwinCell identified candidates not yet approved for SLE but supported by emerging clinical evidence. The model prioritises IL23R and reconstructs causal paths linking it through JAK/STAT intermediaries to DEGs including TNFRSF13B (BAFF), a cytokine known to be involved in the disease (26). This prediction is independently supported by the efficacy of ustekinumab, an IL-12/23 inhibitor already approved for psoriasis and Crohn’s disease, in a phase 2 SLE trial (30). By exploiting interpretable causal evidence supporting the target ranking, TwinCell provides a mechanistic rationale supporting each prediction, enabling researchers to assess biological plausibility before committing to experimental validation.

### TwinBench: A novel benchmarking framework for virtual cells

Rigorous benchmarks are the foundation of progress in any machine learning domain, and the virtual cell field currently needs discriminative evaluation frameworks that rank methods according to their performance on downstream tasks such as target identification. TwinBench addresses this gap by recasting virtual cell evaluation as a recommendation problem, directly measuring whether a model can prioritise the correct intervention among thousands of candidates. The framework introduces two key innovations: first, it uses the rank of the true target as the evaluation metric, replacing transcriptomic similarity scores. Second, it replaces the fixed cutoff *K* of standard recommendation systems with an empirical *p*-value derived from a permutation test, which assesses whether each prediction depends on the biological input signal rather than on patterns memorised from the training set. This corrects for mode collapse and popularity bias, two well-characterised failure modes in which a model systematically favours frequently observed targets regardless of input. We will release TwinBench as an open-source package to accelerate progress toward reliable virtual cell models.

### Perspectives

TwinCell demonstrates competitive generalisation performance across diverse scenarios, ranging from zero-shot *in vitro* cell lines to patient-derived disease states. Furthermore, its ability to generalise to targets absent from the training set highlights its potential for *de novo* target identification. Underlying the predictions, the mechanistic insights provided by the causal graphs offer better confidence and facilitate the exploration of alternative pathway hypotheses. The architecture is highly scalable, generating predictions in seconds and enabling deployment across a broad spectrum of applications, including large scale screening and integration into agentic discovery pipelines.

The model presented in this work was trained on a single perturbation dataset (Tahoe-100M). By extending to additional perturbation datasets we aim to improve performance across therapeutic areas and targets. Future iterations will also integrate higher-quality interactomes, fine-tuned embeddings, cell-cell communication, and multi-species data.

Although virtual cell models have yet to achieve the transformative performance seen in domains such as text and images (14), we believe they are nearing a similar inflection point. Looking ahead, we envision a lab-in-the-loop paradigm where TwinCell generates testable hypotheses that are iteratively refined through experimental feedback, building high-confidence causal cell models across therapeutic areas and positioning virtual cells as translational tools bridging high-throughput *in vitro* experiments and clinical insights.

## Code availability

A waiting list for free access to the TwinCell API and web interface is available on the DeepLife website (www.deeplife.co). The TwinBench package will be provided on GitHub soon.

## Acknowledgement

We thank Moritz Thomas, Christian Tendeng, Sean Gorman, Mayar Ali, Nathalie Jaure and Jonathan Baptista for their critical feedback during the writing of this manuscript. We thank Axel Kuehn and Maxime Gendre for their work on the infrastructure and TwinCell API.

## Material & Methods

### TwinCell LCCM Learning Objective

TwinCell is trained to identify the target *t*^∗^ that maximises the probability given the observed differentially expressed genes DEGs and cell state context **x**:

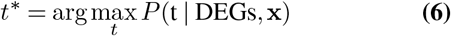

where **x** = (**x**_1_, **x**_2_) are the cell state embeddings obtained from Geneformer foundation model (10), *t* is the target, and DEGs are the differentially expressed genes.

Under a uniform prior over candidate targets, *P* (*t*) = 1*/G* where *G* is the number of proteins in the interactome, and assuming that signalling paths to different DEGs propagate independently through the interactome, the target posterior decomposes as:

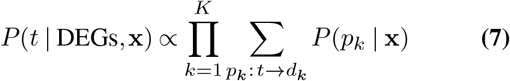

where each *P* (*p*_*k*_ | **x**) is the probability of a signalling path from candidate target *t* to DEG *d*_*k*_. Under first-order Markov signal propagation, each path probability factorises as a product of successive transitions:

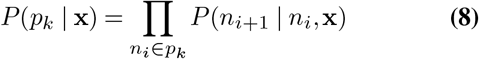

with maximum path length fixed by the interactome diameter (*L* = 7). The model learns cell-state-specific transition probabilities:

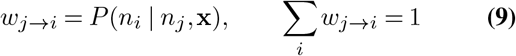

ensuring conservation of probability mass. Edge directionality is enforced by a DAG constraint during training, which ensures acyclicity and a consistent causal ordering from upstream regulators to downstream effectors.

The transition probabilities are optimised end-to-end to maximise *P* (*t* | DEGs, **x**) for the true perturbation target. Once trained, the model enables probabilistic inference for:

1. **Target prioritisation**: Computing *P* (*t* | DEGs, **x**) for all candidate targets and ranking them.
2. **Causal graph construction**: Identifying the signalling routes that maximise 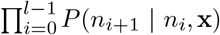 between the DEGs and a selected target, extracting the most probable causal paths.
3. **Downstream effect prediction**: Applying Bayes’ rule to invert the learned conditionals, computing *P* (*d*_*k*_ | *t*, **x**) to identify the impact of a target *t* on a DEG *d*_*k*_.

### TwinBench Metric Details

A model that overfits to the training distribution while ignoring its input (19, 31) exhibits what is termed mode collapse in generative models (32) and popularity bias in recommendation systems (33), analogous to a classifier that always predicts the majority class. In virtual cell modelling, this means a model can predict a mean transcriptomic state while still achieving high *R*^2^ or cosine similarity, because high-dimensional expression profiles share substantial baseline correlation (7, 33, 34).

To address this, TwinBench introduces a two-stage evaluation pipeline (Figure 1b–c): first, scoring and significance testing at the individual target level (Steps 1–2); second, aggregation and metric computation across samples (Steps 3–4). The workflow proceeds as follows.

#### Step 1: Score and Rank Computation

For each experiment (a pair of control and perturbed cell states), the model produces a score for every candidate target (Figure 1b). From these scores, a normalised rank *r*_*j*_ ∈ [0, 1] is computed for each target based on its position relative to all candidates, where *r*_*j*_ = 0 indicates the top-ranked target and *r*_*j*_ = 1 the lowest. The rank of the true perturbation target among all candidates quantifies the model’s ability to prioritise the correct target.

#### Step 2: Empirical p-value Computation

To assess whether predictions depend on the specific biological signal rather than learned priors, we compute an empirical *p*-value for each target via a permutation test (Figure 1c). We perform *N*_*R*_ random permutations of the input differential expression signature. This permutation destroys the gene–gene correlation structure while preserving the distribution of expression values. For each target *t*_*j*_, we compute an empirical *p*-value:

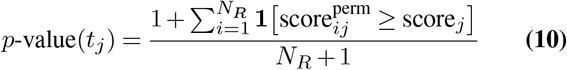

If a target receives a high score even under permuted conditions, the model is driven by the target’s global popularity rather than the specific causal evidence in the query. Conversely, a significant *p*-value indicates the model is actively using the input signal to deviate from its background prior. Targets not expressed in either cell state are excluded from the evaluation, ensuring that unexpressed genes do not inflate the score.

#### Step 3: Building the Target Distribution Across Samples

Steps 1 and 2 produce, for each true target, a normalised rank and an empirical *p*-value. These individual target-level results must be assembled into a joint distribution of ranks and *p*-values before computing summary metrics. The assembly strategy depends on the evaluation setting:

In the *in vitro* setting, multiple samples (cell line– perturbation pairs) each contribute multiple targets. We aggregate them using a two-level bootstrap (*B* = 300): (i) for each experiment *S*_*i*_ with *m*_*i*_ targets, uniformly sample one target to yield a tuple (*r*_*i*_, *p*_*i*_); (ii) draw *n* experiments with replacement; (iii) compute the summary metrics (see Step 4) on the resampled target distribution. This ensures each experiment contributes equally regardless of its target count (35, 36).

In the *in clinico* setting, a single sample (one disease–cell type pair) contributes multiple approved targets. A standard bootstrap (*B* = 300) suffices: draw *m*_*i*_ targets with replacement and compute the metrics on the resampled set (37, 38). In both cases, the bootstrap provides a distribution over the resulting metrics, from which we derive 95% confidence intervals.

#### Step 4: Metrics computation

For each bootstrapped target distribution from Step 3, we compute the Recall, IMNR and F1 score across *p*-value thresholds. Given a threshold *pv*, let *N* (*pv*) denote the number of true targets with *p*-value below *pv*:

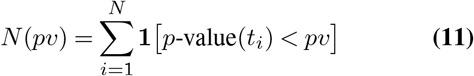

where *N* is the total number of true targets. We then define:

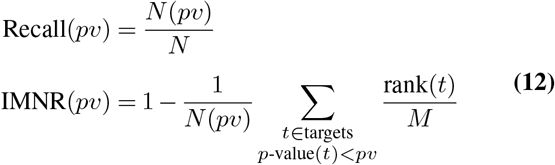

with *M* the number of genes expressed in at least one of the two cell states. Recall measures the fraction of true targets reaching significance, while IMNR measures how highly ranked the significant targets are (IMNR = 1 if all significant targets are top-ranked, IMNR = 0.5 at random). Combining both into the F1 score:

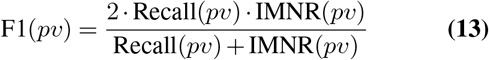

We evaluate across 30 linearly-spaced *p*-value thresholds in [0.01, 1.0] and compute the Area Under the Curve (AUC) of the F1 score. For a random predictor with uniform *p*-values, recall at threshold *pv* equals *pv* and IMNR equals 0.5, yielding a theoretical random baseline of AUC-F1_random_ = 45%. To aid interpretation, we normalise such that the AUC-F1 falls in [0, 1] with 0.5 corresponding to random performance:

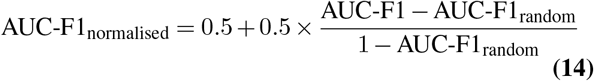

The bootstrap from Step 3 provides a distribution of AUC-F1 values, from which we report the median and 95% confidence interval.

### TwinBench on Generative Models

TwinBench can be readily adapted to evaluate conditional generative models such as STATE (13), where the objective is to predict a final cellular state **x**_2_ given an initial state **x**_1_ and a perturbation. We decompose such models into an embedding function *f*_emb_ and a transition function *f*_trans_. Given input **x**_1_ and ground-truth output **x**_2_, we compute:

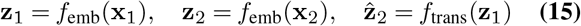

The evaluation metric is the cosine similarity between predicted and true states:

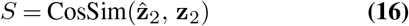

For the permutation test, we randomly permute the features of **x**_1_ to obtain 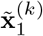 and compute the corresponding prediction:

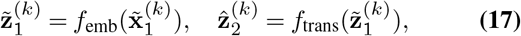

and similarity score:

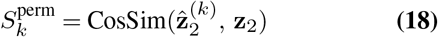

The empirical *p*-value is then:

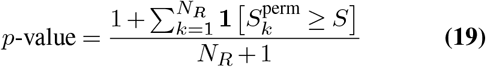

A non-significant *p*-value indicates that the model’s prediction is independent of the gene–gene correlation structure in **x**_1_, diagnosing mode collapse: the model produces a generic average response rather than a perturbation-specific prediction.

### Generalisation scenarios

#### In vitro

The *in vitro* evaluation uses a train/test split constructed along two zero-shot axes: held-out cell lines and held-out perturbations. The held-out cell lines consist of the original STATE hold-out set (13), supplemented with additional cell lines chosen so that liver and kidney tissues are entirely absent from training, thereby introducing tissue-level out-of-distribution contexts into the test set (Table S1). The held-out perturbations exclude not only individual drugs but also their complete target sets from the training data (Table S2). The combined held-out set is the union of these two axes, yielding a 20% test split.

#### In clinic

The *in clinico* scenario benchmarks model predictions against clinically validated drug–target–disease associations curated in the Open Targets Platform (version 25.06) (39). We extracted the protein targets of compounds that have reached Phase III or Phase IV clinical status across five therapeutic areas: Systemic Lupus Erythematosus, Parkinson Disease, Psoriasis, Crohn’s Disease, and Ulcerative Colitis (Table S3).

### Multiomics Interactome

We used a proprietary interactome comprising 15K nodes and 201K edges, including protein–protein and transcriptional interactions. The interactome was built by aggregating and harmonising protein and gene interaction data from close to a hundred databases including BioGRID (40), Reactome (41), and IntAct (42). Only experimentally determined entries were considered, excluding computationally inferred ones. Interactions were restricted to direct physical connections rather than those potentially mediated by unknown intermediary molecules. Recommended filtering for high confidence was applied for each database when available. We included only interactions found in at least two independent databases, unless they originated from a single database with high standards of manual curation, such as SIGNOR (43).

### Linear Ridge Regression as a baseline for recommendation systems

To evaluate the comparative performance of our proposed models, we employ Linear Ridge Regression as a robust baseline Figure S2. In the context of collaborative filtering, we model the recommendation problem as an item-item regression task, where the goal is to learn a weight matrix *W* that reconstructs the user-item interaction matrix *X*. Because user-item interaction data is inherently sparse and high-dimensional, standard least squares estimation is prone to overfitting and numerical instability. Ridge regression addresses this by introducing an *L*_2_ regularization term to the objective function:

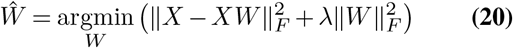

where ∥ · ∥_*F*_ denotes the Frobenius norm and *λ >* 0 is a hyperparameter controlling the regularization strength. The constraint *diag*(*W*) = 0 is typically applied to prevent the model from learning identity mappings. Solving this optimisation problem via its closed-form analytical solution allows for computationally efficient training, providing a deterministic and competitive baseline consistent with established linear methods such as EASE (19).

To assess the impact of input representation on linear baseline performance, we evaluated two distinct preprocessing strategies:

1. **Geneformer embeddings**: Individual single-cell RNA-seq profiles were embedded using the pre-trained Geneformer foundation model, then averaged per condition to produce a single fixed-dimensional vector per cell state.
2. **Highly Variable Genes (HVG)**: The top 5,000 highly variable genes were selected across the training set using standard variance-based filtering. The samples are then pseudobulked and normalised using log1p.

For the TwinBench permutation test, each preprocessing strategy has its own randomisation procedure: for Gene-former embeddings, the input gene expression is permuted before applying the foundation model, as described in the generative model adaptation above; for HVG, the selected highly variable gene values are permuted directly.

### Network Medicine Baseline

Network Medicine relies on computing the topological distance between observed DEGs and drugs modules defined by their targets within the interactome (20). Unlike the other models that learn to predict targets directly, this method calculates a distance score assessing whether drug targets are closer to the disease module (DEGs) than expected by chance in the interaction graph. Because this approach inherently includes a permutation test using degree preservation to compute its specific *z*-score and *p*-value to compare the observed distance to a distribution of distances generated at random, we used these pre-computed metrics directly within the TwinBench framework.

### Omics Dataset Description

#### In vitro dataset

The *Tahoe-100M* atlas represents a giga-scale single-cell perturbation resource comprising 100 million transcriptomic profiles measuring 191 small-molecule perturbations on 50 cancer cell lines (12).

#### In in clinico dataset

We built atlases using our internal pipeline on psoriasis, ulcerative colitis, Crohn and Parkinson disease. Regarding the systemic lupus erythematosus (SLE) atlas, we selected dataset GSE174188 (21), which comprises over 1.2 million cells profiled from 161 SLE patients and 99 healthy controls.

Raw, annotated count matrices were obtained from the cellxgene portal. scRNA-seq data was reprocessed using DeepLife’s pipeline, including cell- and gene-level quality control. Briefly, cells were filtered based on total UMI counts (maximum of 8,000), number of detected genes (minimum of 300 and maximum of 2,200), doublet probability estimated using Scrublet (44), and mitochondrial RNA content, with cells exceeding 20% mitochondrial reads excluded as likely stressed or dying cells. Highly variable genes were identified using Scanpy’s `pp.highly_variable_genes` function, the top 4,000 highly variable genes were retained for downstream analyses. Dimensionality reduction was performed by principal component analysis (PCA) on the highly variable genes, computing the first 15 principal components using `pp.pca`. To mitigate technical batch effects across samples, Harmony (45) was applied to the PCA embeddings. A neighbourhood graph was then constructed on the first 50 Harmony-adjusted principal components using `pp.neighbours` with 15 nearest neighbours. For two-dimensional visualisation, the neighbourhood graph was embedded using UMAP via `tl.umap`, with a minimum distance parameter set to 0.5. Cell clustering was performed using the Leiden algorithm on the integrated neighbourhood graph. To improve cellular resolution, selected clusters were iteratively re-clustered at varying resolution parameters, enabling finer subdivision of heterogeneous populations. Cluster identities were assigned through manual annotation based on marker genes reported in the literature, complemented by automated cell type labels generated using Pan-Azimuth. All preprocessing and analyses steps of scRNA-seq data were run in python 3 using Scanpy (46) and anndata (47) unless otherwise stated.

## Supplementary Materials

**Fig. S1.**
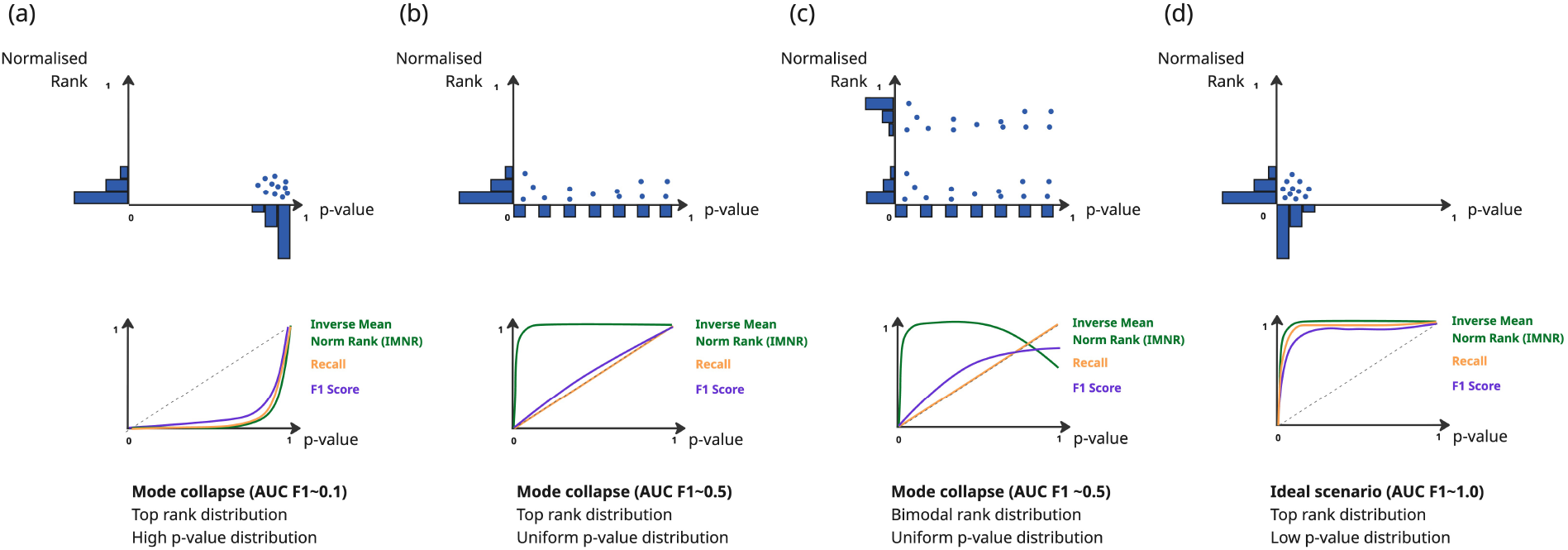
TwinBench diagnostic scenarios illustrating common metric patterns. Each column represents a distinct model behaviour, characterised by the joint distribution of empirical *p*-values and normalised ranks, and the resulting IMNR, Recall and F1 curves across *p*-value thresholds. (a) Mode collapse with high *p*-values (AUC F1∼0.1): true targets are top-ranked but their scores are not significantly higher than those obtained from permuted inputs, indicating that the ranking reflects learned popularity rather than input-specific signal. (b) Mode collapse with uniform *p*-values (AUC F1∼0.5): true targets are again top-ranked, but *p*-values are uniformly distributed, meaning the model cannot distinguish real biological signal from noise; Recall grows linearly with the *p*-value threshold, matching random expectation. (c) Mode collapse with bimodal ranks and uniform *p*-values (AUC F1∼0.5): a fraction of targets are well-ranked while others are not, and the uniform *p*-value distribution again indicates input-insensitive predictions. (d) Ideal scenario (AUC F1∼1.0): true targets are both top-ranked and achieve low *p*-values, confirming that the model leverages the specific biological structure of the input; IMNR, Recall and F1 all rise steeply at stringent *p*-value thresholds.

**Fig. S2.**
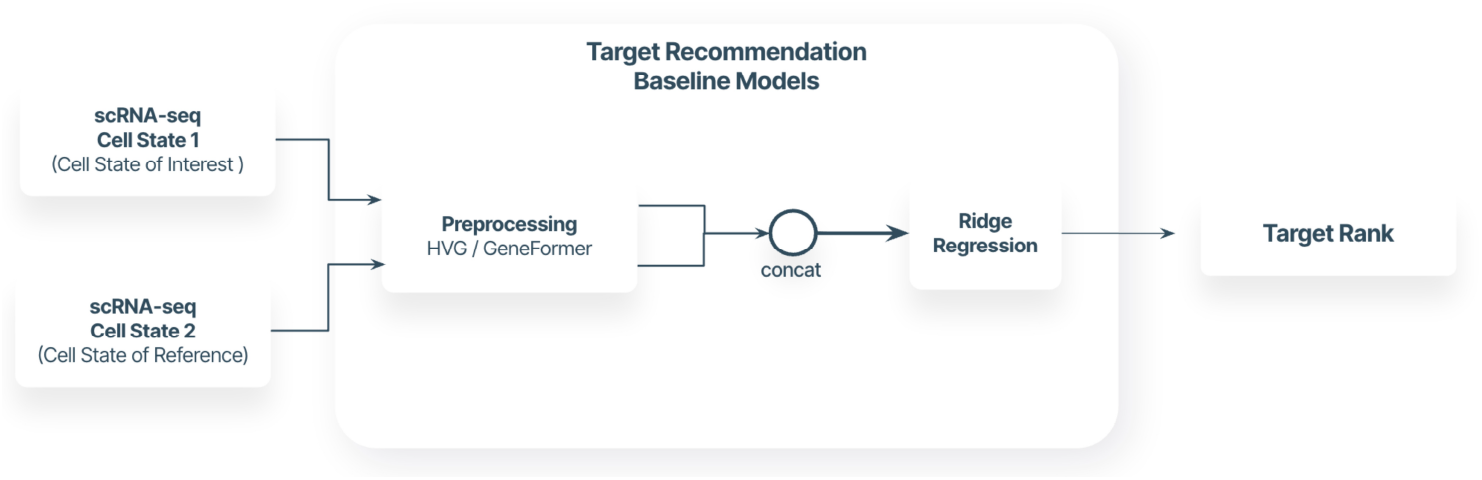
Linear Ridge Regression as a baseline for recommendation systems. Two input cell states are processed using various feature-extraction methods: selection of highly variable genes (HVGs) and Geneformer embeddings on pseudobulk profiles. The processed features are concatenated and fed into a Ridge regression model trained to predict target scores, adapting the EASE architecture (19) to this domain.

**Table S1.** Held-out cell lines in the *in vitro* test set. Cell lines are listed with their tissue of origin

| Cell line | Organ |
| --- | --- |
| CVCL_0480 | Pancreas |
| CVCL_1056 | Kidney |
| CVCL_0366 | Liver |
| CVCL_1285 | Lung |
| CVCL_1693 | Lung |
| CVCL_0334 | Pancreas |
| CVCL_1097 | Skin |
| CVCL_1531 | Lung |
| CVCL_1098 | Liver |

**Table S2.**
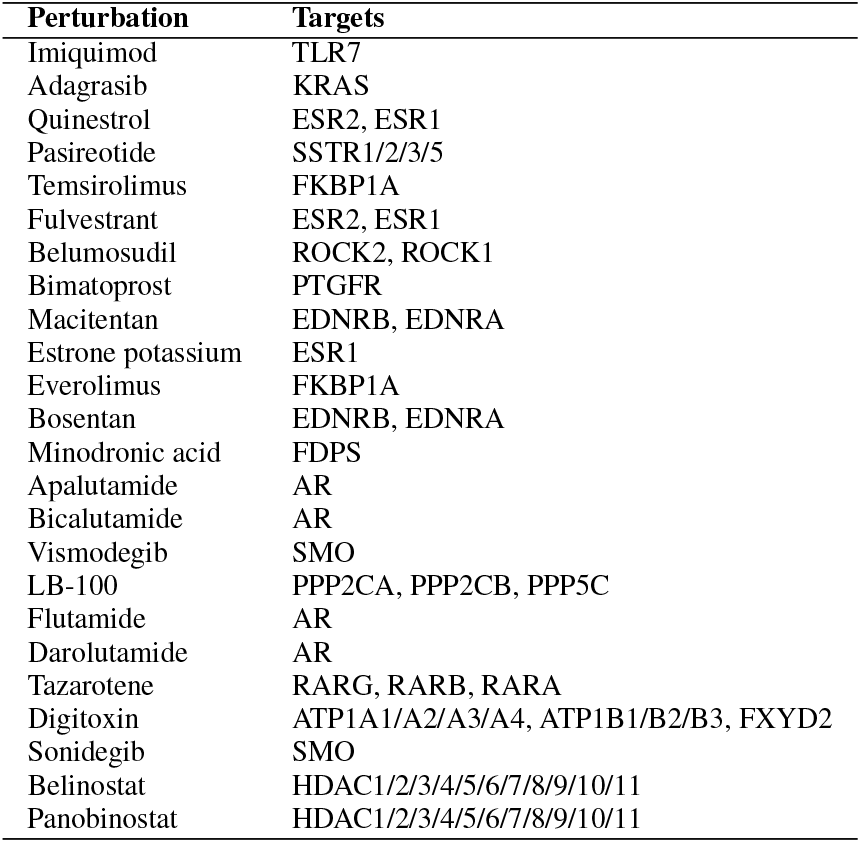
Held-out perturbations in the *in vitro* test set. Perturbations are listed with their molecular targets associated using Open Target (version 25.06)

**Table S3.**
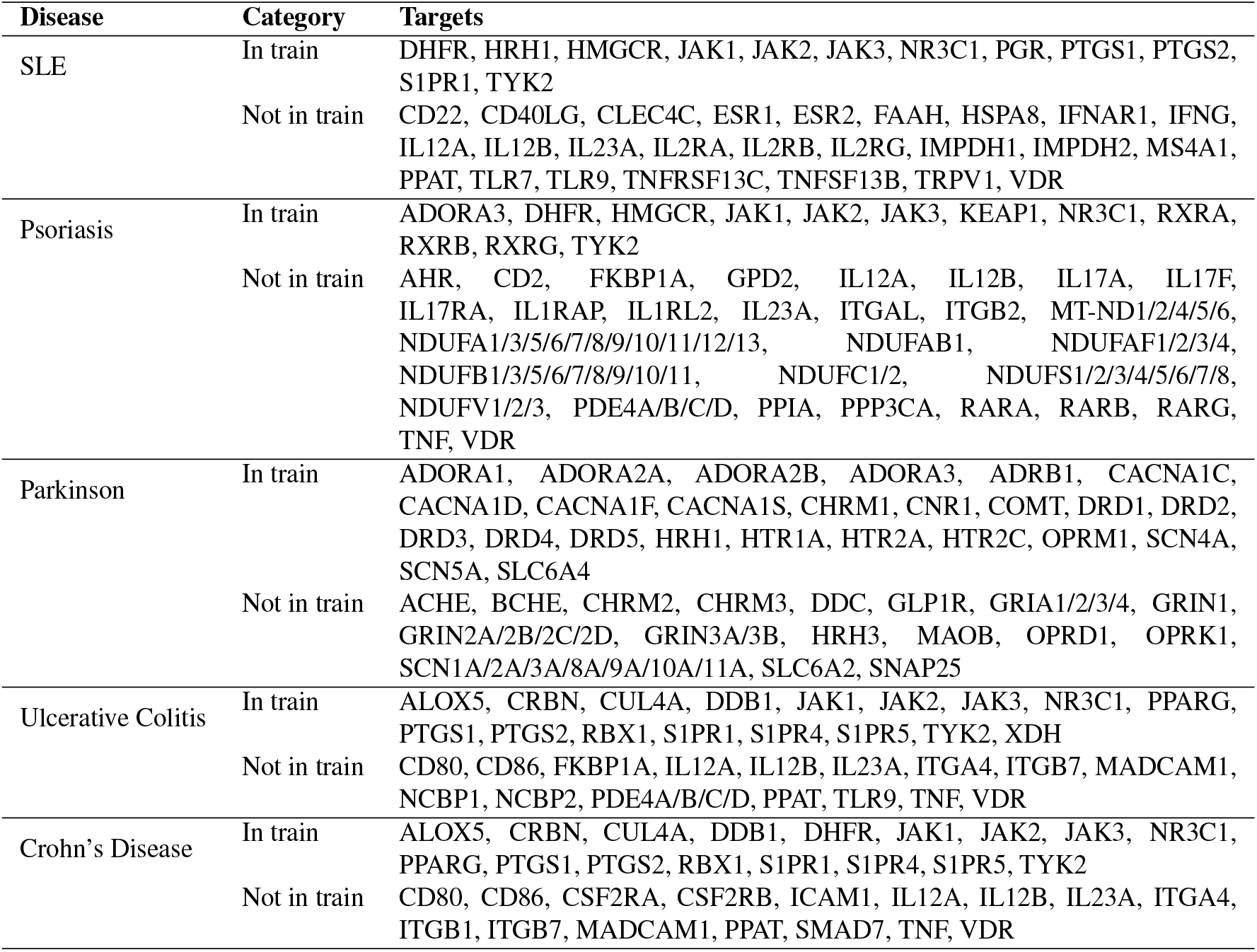
Approved drug targets per disease used in *in clinico* benchmarking. Targets are categorised by their presence in the Tahoe 100M *in vitro* training set.

| Disease | Category | Targets |
| --- | --- | --- |
| SLE | In train | DHFR, HRH1, HMGCR, JAK1, JAK2, JAK3, NR3C1, PGR, PTGS1, PTGS2, S1PR1, TYK2 |
|  | Not in train | CD22, CD40LG, CLEC4C, ESR1, ESR2, FAAH, HSPA8, IFNAR1, IFNG, IL12A, IL12B, IL23A, IL2RA, IL2RB, IL2RG, IMPDH1, IMPDH2, MS4A1, PPAT, TLR7, TLR9, TNFRSF13C, TNFSF13B, TRPV1, VDR |
| Psoriasis | In train | ADORA3, DHFR, HMGCR, JAK1, JAK2, JAK3, KEAP1, NR3C1, RXRA, RXRB, RXRG, TYK2 |
|  | Not in train | AHR, CD2, FKBP1A, GPD2, IL12A, IL12B, IL17A, IL17F, IL17RA, IL1RAP, IL1RL2, IL23A, ITGAL, ITGB2, MT-ND1/2/4/5/6, NDUFA1/3/5/6/7/8/9/10/11/12/13, NDUFAB1, NDUFAF1/2/3/4, NDUFB1/3/5/6/7/8/9/10/11, NDUFC1/2, NDUFS1/2/3/4/5/6/7/8, NDUFV1/2/3, PDE4A/B/C/D, PPIA, PPP3CA, RARA, RARB, RARG, TNF, VDR |
| Parkinson | In train | ADORA1, ADORA2A, ADORA2B, ADORA3, ADRB1, CACNA1C, CACNA1D, CACNA1F, CACNA1S, CHRM1, CNR1, COMT, DRD1, DRD2, DRD3, DRD4, DRD5, HRH1, HTR1A, HTR2A, HTR2C, OPRM1, SCN4A, SCN5A, SLC6A4 |
|  | Not in train | ACHE, BCHE, CHRM2, CHRM3, DDC, GLP1R, GRIA1/2/3/4, GRIN1, GRIN2A/2B/2C/2D, GRIN3A/3B, HRH3, MAOB, OPRD1, OPRK1, SCN1A/2A/3A/8A/9A/10A/11A, SLC6A2, SNAP25 |
| Ulcerative Colitis | In train | ALOX5, CRBN, CUL4A, DDB1, JAK1, JAK2, JAK3, NR3C1, PPARG, PTGS1, PTGS2, RBX1, S1PR1, S1PR4, S1PR5, TYK2, XDH |
|  | Not in train | CD80, CD86, FKBP1A, IL12A, IL12B, IL23A, ITGA4, ITGB7, MADCAM1, NCBP1, NCBP2, PDE4A/B/C/D, PPAT, TLR9, TNF, VDR |
| Crohn's Disease | In train | ALOX5, CRBN, CUL4A, DDB1, DHFR, JAK1, JAK2, JAK3, NR3C1, PPARG, PTGS1, PTGS2, RBX1, S1PR1, S1PR4, S1PR5, TYK2 |
|  | Not in train | CD80, CD86, CSF2RA, CSF2RB, ICAM1, IL12A, IL12B, IL23A, ITGA4, ITGB1, ITGB7, MADCAM1, PPAT, SMAD7, TNF, VDR |

**Table S4.** Top 30 predicted proteins from TwinCell analysis of SLE CD4^+^ activated T cells. ranked by the TwinCell score. Score: log-unnormalised posterior log *P* (*t* | DEGs, x) up to an additive constant shared across all targets; Rank: absolute rank among all interactome proteins; Rank %: percentile rank in the interactome; *z*-score: standardised effect size relative to permuted scores; Approved targets from OpenTargets (phase III/IV) are indicated with ✓.

| Gene | Score | Rank | Rank (%) | z-score | Approved Target |
| --- | --- | --- | --- | --- | --- |
| STAT2 | 212.87 | 0 | 0.000 | 18.40 |  |
| IRF9 | 207.25 | 1 | 0.005 | 43.26 |  |
| STAT1 | 203.94 | 2 | 0.010 | 8.32 |  |
| IFNAR2 | 202.46 | 3 | 0.015 | 14.73 |  |
| CD55 | 199.47 | 4 | 0.020 | 9.98 |  |
| CREBBP | 188.66 | 5 | 0.025 | 4.81 |  |
| CD14 | 177.45 | 6 | 0.029 | 6.66 |  |
| EP300 | 169.83 | 7 | 0.034 | 4.07 |  |
| EIF2AK2 | 161.24 | 8 | 0.039 | 9.31 |  |
| TYK2 | 160.96 | 9 | 0.044 | 5.15 | ✓ |
| NUCB1 | 153.17 | 10 | 0.049 | 2.97 |  |
| PEX19 | 150.10 | 11 | 0.054 | 6.25 |  |
| STAT6 | 149.63 | 12 | 0.059 | 8.56 |  |
| FGFR1 | 144.76 | 13 | 0.064 | 3.28 |  |
| PTGS2 | 142.13 | 14 | 0.069 | 3.21 | ✓ |
| STAT5A | 141.04 | 15 | 0.074 | 4.37 |  |
| OSM | 140.06 | 16 | 0.078 | 6.78 |  |
| ZNF346 | 138.93 | 17 | 0.083 | 6.64 |  |
| EEF1E1 | 138.41 | 18 | 0.088 | 2.00 |  |
| IFT20 | 137.17 | 19 | 0.093 | 10.02 |  |
| STAT3 | 135.52 | 20 | 0.098 | 2.94 |  |
| IL23R | 135.49 | 21 | 0.103 | 5.46 |  |
| ECH1 | 134.00 | 22 | 0.108 | 5.23 |  |
| LTB | 132.43 | 23 | 0.113 | 6.74 |  |
| TRAF3IP1 | 131.46 | 24 | 0.118 | 9.64 |  |
| ISG15 | 130.76 | 25 | 0.123 | 5.94 |  |
| TLR3 | 130.33 | 26 | 0.128 | 5.65 |  |
| DRAP1 | 128.48 | 27 | 0.132 | 3.42 |  |
| FANCG | 128.41 | 28 | 0.137 | 3.96 |  |
| NR3C1 | 128.08 | 29 | 0.142 | 2.30 | ✓ |

**Table S5.** Open Targets approved drug targets in SLE ranked by TwinCell analysis of CD4^+^ activated T cells. This table shows the ranking of clinically validated SLE targets expressed in CD4^+^ activated T cells (from Open Targets, phase III/IV clinical trials) according to TwinCell’s target prioritisation score. Score: TwinCell target score; Rank %: percentile rank in the interactome; *z*-score: standardised effect size;

| Gene | Score | Rank | Rank (%) | z-score |
| --- | --- | --- | --- | --- |
| TYK2 | 160.96 | 9 | 0.088 | 5.14 |
| PTGS2 | 142.13 | 14 | 0.137 | 3.08 |
| NR3C1 | 128.08 | 29 | 0.285 | 2.26 |
| IL23A | 121.18 | 37 | 0.363 | 5.85 |
| JAK3 | 114.22 | 54 | 0.530 | 4.34 |
| JAK2 | 113.72 | 55 | 0.540 | 2.62 |
| JAK1 | 96.77 | 121 | 1.187 | 3.27 |
| DHFR | 87.60 | 181 | 1.776 | 2.11 |
| IL12A | 75.38 | 292 | 2.865 | 2.67 |
| ESR1 | 68.86 | 366 | 3.591 | 1.17 |
| IMPDH2 | 67.60 | 378 | 3.708 | 2.44 |
| IFNAR1 | 63.36 | 435 | 4.268 | 8.20 |
| ESR2 | 32.78 | 1139 | 11.174 | 2.14 |
| IL2RB | 31.30 | 1198 | 11.753 | 3.85 |
| IL2RG | 31.11 | 1206 | 11.832 | 3.52 |
| HSPA8 | 24.24 | 1526 | 14.971 | 1.47 |
| VDR | 20.04 | 1782 | 17.483 | 0.57 |
| HMGCR | 13.11 | 2321 | 22.771 | −0.22 |
| IFNG | 10.56 | 2578 | 25.292 | 1.57 |
| IL2RA | 4.18 | 3742 | 36.711 | 1.66 |
| CD22 | 3.37 | 3910 | 38.360 | 0.35 |
| TNFSF13B | 2.69 | 4086 | 40.086 | −0.96 |
| TLR7 | 2.64 | 4094 | 40.165 | 0.58 |
| S1PR1 | 1.82 | 4373 | 42.902 | −0.50 |
| CD40LG | 0.59 | 5028 | 49.328 | 0.02 |
| MS4A1 | 0.12 | 5638 | 55.312 | 0.15 |
| TNFRSF13C | 0.07 | 5774 | 56.647 | 0.02 |

**Table S6.** Top 5 paths from IL23R to TNFRSF13B (BAFF) in SLE CD4^+^ activated T cells. Each row represents a signalling path inferred, with the associated probability. Paths are ordered by decreasing probability.

| Path | Probability |
| --- | --- |
| IL23R → TYK2 → STAT1 → TNFRSF13B | $2.00 \times 10^{-7}$ |
| IL23R → STAT3 → STAT1 → TNFRSF13B | $4.02 \times 10^{-11}$ |
| IL23R → JAK2 → STAT1 → TNFRSF13B | $1.46 \times 10^{-11}$ |
| IL23R → TYK2 → STAT1 → EP300 → TNFRSF13B | $1.35 \times 10^{-11}$ |
| IL23R → TYK2 → STAT5A → STAT1 → TNFRSF13B | $8.99 \times 10^{-12}$ |

